# A healthy childhood environment helps to combat inherited susceptibility to obesity

**DOI:** 10.1101/2020.01.13.905125

**Authors:** Anke Hüls, Marvin N Wright, Leonie H Bogl, Jaakko Kaprio, Lauren Lissner, Denes Molnár, Luis Moreno, Stefaan De Henauw, Alfonso Siani, Toomas Veidebaum, Wolfgang Ahrens, Iris Pigeot, Ronja Foraita, on behalf of the IDEFICS/I.Family consortia

## Abstract

**Objectives:** To investigate the degree by which the inherited susceptibility to obesity is modified by environmental factors during childhood and adolescence.

**Design:** Cohort study with repeated measurements of diet, lifestyle factors and anthropometry.

**Setting:** The pan-European IDEFICS/I.Family cohort

**Participants:** 8,609 repeated observations from 3,098 children aged 2 to 16 years, examined between 2007 and 2014.

**Main outcome measures:** Body mass index (BMI) and waist circumference. Genome-wide polygenic risk scores (PRS) to capture the inherited susceptibility of obesity were calculated using summary statistics from independent genome-wide association studies of BMI. Gene-environment interactions of the PRS with sociodemographic (European region, socioeconomic status) and lifestyle factors (diet, screen time, physical activity) were estimated.

**Results:** The PRS was strongly associated with BMI (r^2^ = 0.11, p-value = 7.9 × 10^−81^) and waist circumference (r^2^ = 0.09, p-value = 1.8 × 10^−71^) in our cohort. The associations with BMI increased from r^2^=0.03 in 3-year olds to r^2^=0.18 in 14-year olds and associations with waist circumference from r^2^=0.03 to r^2^=0.14. Being in the top decile of the PRS distribution was associated with 3.63 times higher odds for obesity (95% confidence interval (CI): [2.57, 5.14]). We observed significant interactions with demographic and lifestyle factors for BMI as well as waist circumference. The risk of becoming obese among those with higher genetic susceptibility was ~38% higher in children from Southern Europe (BMI: p-interaction = 0.0066, Central vs. Southern Europe) and ~61% higher in children with a low parental education (BMI: p-interaction = 0.0012, low vs. high). Furthermore, the risk was attenuated by a higher intake of dietary fiber (BMI: p-interaction=0.0082) and shorter screen times (BMI: p-interaction=0.018).

**Conclusions:** Our results highlight that a healthy childhood environment might partly offset a genetic predisposition to obesity during childhood and adolescence.

## Introduction

Obesity is a complex multifaceted condition and its prevalence has been increasing continuously over previous decades and has reached a high plateau in Western countries [1]. In 2015, a total of 107.7 million children and 603.7 million adults were obese. Although the prevalence of obesity among children has been lower than that among adults, the rate of increase in childhood obesity has been greater than the rate of increase in adult obesity, which is most likely due to adverse changes of environmental and demographic factors with a direct impact on children’s health [2].

With the advent of genome-wide association studies (GWAS), it was shown that multiple genetic loci increase the susceptibility to obesity [3,4]. However, genome-wide significant variants identified in the first large-scale GWAS on body mass index (BMI) only account for a small portion of BMI variation (~2.7%) [3]. A more recent genome-wide meta-analysis extended the number of individuals from ~300,000 [3] to ∼700,000 [4], which consequently increased the number of genome-wide significant SNPs from 97 to 751. Even these 751 genome-wide significant SNPs account for only ∼6.0% of the variance of BMI [4]. However, genome-wide estimates suggest that common variation accounts for >20% of BMI variation [3], which highlights the polygenic architecture of BMI. More recently, whole genome data even increased the fraction of variance of BMI accounted for by genetic variants, both common and rare, to 40% [5]. From twin studies we know that the heritability of BMI also depends on socioeconomic status [6] and physical activity [7], suggesting that when socioeconomic status or physical activity is high, genetic factors become less influential. Using candidate SNPs - either single genotypes or <100 SNPs combined in a polygenic risk score (PRS), which is defined as a weighted sum of BMI-related risk alleles - it was further shown that the genetic predisposition to obesity is attenuated by a healthy lifestyle including physical activity [8,9] and adherence to healthy dietary patterns [9–15]. However, most previous gene-environment (GxE) interaction studies primarily involved adults [8–15] or used only a candidate SNP [16], so that it is unknown whether the inherited susceptibility to obesity is modified by environmental factors already during childhood and adolescence. Another limitation of previous gene-environment interaction analyses is that they were based on <100 SNPs that reached genome-wide significance in previous GWAS on BMI [3], which do not capture the whole polygenic risk profile of obesity due to their low heritability. Khera et al. suggested that the power to predict BMI by PRS can be improved by using lower p-value thresholds or even genome-wide approaches [17]. Using a genome-wide polygenic risk score based on effect estimates from [3], Khera et al. reported that the PRS-effect on weight and BMI z-scores emerges early in life and increases until adulthood and that a high PRS is a strong risk factor for severe obesity and associated diseases [17]. The authors suggested that given that the weight trajectories of individuals in different PRS deciles start to diverge early in childhood, targeted strategies for obesity prevention may have maximal effect when employed early in life. However, because lifestyle factors were not considered in their study, it is not known to which degree the genetic predisposition to obesity is modifiable by a healthy lifestyle early in life. Another limitation of [17] is the use of weight and BMI as only proxies for obesity. Since several studies have shown that classifying obesity using BMI alone misses an increasing proportion of individuals categorized as obese [18,19], it is important to test the performance of BMI-PRS for the prediction of waist circumference, which is proposed to be a better proxy for obesity-associated metabolic abnormalities [20].

In this study, 1) we show the prediction capacity of the PRS proposed in [17] for BMI as well as for waist circumference of European children and adolescents and 2) analyze its interaction with parental education, region of residence, selected dietary variables and physical activity to investigate to which degree the inherited susceptibility to obesity in children is modified by these sociodemographic and lifestyle factors. The analyses are based on 8,609 repeated observations from 3,098 children and adolescents aged 2 to 16 years from the pan-European IDEFICS/I.Family cohort.

## Methods

### Study Population

The pan-European IDEFICS/I.Family cohort [21,22] is a multi-center, prospective study on the association of social, environmental and behavioral factors with children’s health status. Children were recruited through kindergarten or school settings in Belgium, Cyprus, Estonia, Germany, Hungary, Italy, Spain and Sweden. In 2007/2008, 16,229 children aged between 2 and 9.9 years participated in the baseline survey. Follow-up surveys were conducted after two (FU1, N = 11,043 plus 2,543 newcomers) and six years (FU2, N = 7,117 plus 2,512 newly recruited siblings). Physical examinations covered a broad spectrum of parameters according to a detailed and standardized study protocol. Questionnaires were completed by parents for children younger than 12 years. In the second follow-up (FU2), adolescents of 12 years of age or older reported for themselves. All questionnaires were developed in English and translated into local languages. The quality of translations was checked by back translation into English. The study was conducted in agreement with the Declaration of Helsinki; all procedures were approved by the local ethics committees and written and oral informed consents were obtained from the parents, their children and adolescents, respectively, as applicable. Children were selected for a whole-genome scan based on their participation in the individual study modules. Children from Cyprus were not included in this initial genotyping to minimize population stratification.

### Assessment of BMI and Waist Circumference

BMI was calculated as weight divided by height squared [kg/m^2^]. Height was measured to the nearest 0.1 cm by a SECA 225 Stadiometer (Seca GmbH & Co. KG., Hamburg, Germany) and body weight was measured in fasting state in light underwear on a calibrated scale accurate to 0.1 kg by a Tanita BC 420 SMA scale (TANITA, Tokyo, Japan). Waist circumference was measured in upright position with relaxed abdomen and feet together using an inelastic tape (Seca 200, Birmingham, UK), precision 0.1 cm, midway between the iliac crest and the lowest rib margin to the nearest 0.1 cm [23]. Age- and sex-specific BMI and waist circumference z-scores for children and adolescents were calculated using reference data from the International Obesity Task Force [24] and from British children [25], respectively.

### Genotyping and Quality Control

DNA was extracted from saliva or blood samples using established procedures. Genotyping of 3,515 children was performed on the UK Biobank Axiom array (Santa Clara, USA) in two batches (2015 and 2017). Following the recommendations of [26], sample and genotype quality control measures were applied (see supplementary materials for details), resulting in 3,099 children and 3,424,677 genotypes after imputation. A genetic relatedness matrix was calculated to account for the degree of relatedness within the study sample and to adjust for population stratification [27,28] by using the program EMMAX (https://genome.sph.umich.edu/wiki/EMMAX).

### Polygenic Risk Score Calculation

We calculated PRS based on genome-wide summary statistics for BMI from European ancestry populations. The PRS (called PRS-Khera) was proposed in [17]. It consists of 2,100,302 SNPs and is based on summary statistics from the first large-scale GWAS of BMI (~300,000 samples) [3]. PRS-Khera was calculated in [17] using a computational algorithm called LDPred, which is a Bayesian approach to calculate a posterior mean effect for all variants using external weights with subsequent shrinkage based on linkage disequilibrium [29]. Using LDPred, each variant was reweighted according to the prior GWAS [3], the degree of correlation between a variant and others nearby, and a tuning parameter that denotes the proportion of variants with non-zero effect.

In sensitivity analyses, the performance of PRS-Khera was compared to the PRS calculated with PRSice [30] and the PRS based on only genome-wide significant SNPs from two reference populations (same reference population as for PRS-Khera (~300,000 samples) [3] and the largest published GWAS study of BMI to date (~700,000 samples) [4]). More details on the different PRS are given in the supplementary methods and Figures S1 to S3.

### Assessment of Dietary Intake

We used long-term and short-term dietary measurements assessed by food frequency questionnaires (FFQs) and repeated 24 hour dietary recalls, respectively [31]. A fruit and vegetable score was calculated from FFQs (for more details on the FFQs and calculation of the fruit and vegetable score, see supplemental material). We expressed the fruit and vegetable consumption as the relative frequency in relation to all foods reported in the FFQs [32]. The FFQs were self-reported by adolescents 12 years and older and proxy-reported by a parent or other caregiver for children below the age of 12 years.

Energy and dietary fiber intake were assessed by repeated 24 hour dietary recalls [33,34]. Usual intakes for fiber were estimated based on the validated National Cancer Institute (NCI) method, which is one of the most widely accepted methods for this purpose [35,36]. This method allows for the inclusion of covariates such as age and accounts for different intakes on weekend days vs. weekdays, and further corrects for the day-to-day variation in energy and fiber intakes. Usual intakes were estimated for each child stratified by sex and considering age as a covariate. Fiber intake was here expressed in relation to total energy intake in mg/kcal. See supplemental material for more details.

### Assessment of Physical Activity

Physical activity was objectively measured by using Actigraph's uniaxial or three-axial accelerometers [37,38]. At baseline and FU1, children were asked to wear the accelerometer for three days (including one weekend day) and at FU2 for a full week during waking hours (except when swimming or showering). The accelerometers were attached to the right hip with an elastic belt. Participants (either the parents or the adolescents themselves) were given written instructions on how to use the accelerometer and were asked to complete diaries to record non-wear times of the device. The daily average cumulative duration of time spent in moderate-to-vigorous physical activity (MVPA) was expressed as minutes per day according to previously defined cut-off values [39]. Especially for children, accelerometer measurements are far less prone to measurement errors than self-reported activities through questionnaires [40,41]. See supplementary material for more details.

### Assessment of Screen Time

Screen time was assessed by asking how many hours per day the child/adolescent usually spends watching television (including videos or DVDs) and by another question on the time sitting in front of a computer and game console [42,43]. Responses were weighted and summed across weekdays and weekend days and the quantified frequencies from both questions were added to create a continuous variable of total screen time in hours per day. Parents reported for children younger than 12 years, while older children (≥ 12 years) reported for themselves. See supplemental material for more details.

### Assessment of Sociodemographic Variables

Parental education was retrieved from questionnaires and coded according to the International Standard Classification of Education (ISCED) [44]. For the analyses, the highest parental education of both parents was coded as low (ISCED levels 1 and 2; ≤9 years of education), medium (ISCED levels 3 and 4) and high (ISCED levels 5 and 6; ≥2 years of education after high school). The region of residence was coded as Northern Europe (Estonia, Sweden), Central Europe (Belgium, Germany, and Hungary) and Southern Europe (Italy, Spain).

### Statistical Analyses

Our data consist of up to three repeated measurements of individuals, some of which were siblings. We used generalized linear mixed models where the covariance matrix of the random intercept is proportional to a genetic relatedness matrix. We applied the generalized linear mixed model approach of Chen et al. [27] that jointly controls for relatedness and population stratification. All models were adjusted for sex, age, region of residence and parental education. All models that did not include fiber intake were additionally adjusted for the vegetable score. When testing associations with categorical variables (sex, region of residence and parental education), we used the category with the largest sample size as reference category.

All p-values from the gene-environment interaction analyses were adjusted according to the number of tested environmental factors using the false-discovery rate (FDR). We reported 95% confidence intervals (95% CI) and two-sided p-values, and considered p-values less than 0.05 statistically significant. We used R 3.5.1 [45] for all statistical analyses.

## Results

The study sample included 8,609 repeated BMI measurements from at maximum three time points (baseline, FU1, FU2) of 3,098 children aged 2 to 16 years (Table 1). The number of participants decreased only slightly between the follow-up investigations from n = 3,016 at baseline (mean age 6 years) to n = 2,656 at FU2 (mean age 12 years). Half of the children were girls, most children came from families with a medium or high level of education and the majority lived in Central European countries. The distributions of the dietary variables (vegetable score and fiber intake) and time spent in MVPA were similar between baseline and the two follow-up samples, whereas children and adolescents spent more time in front of screens at FU1 and FU2 as compared to baseline. On average, BMI and waist circumference of our analysis group were higher than in the reference populations [24,25] (mean z-scores > 0).

**Table 1.** Study characteristics of the 8,609 repeated observations from 3,098 children.

|  |  | Baseline | First follow-up<br>(FU1) | Second follow-up<br>(FU2) |
| --- | --- | --- | --- | --- |
| **n** |  | 3016 | 2937 | 2656 |
| **Age (years)** |  |  |  |  |
|  | Mean (SD) | 6.19 (1.77) | 8.12 (1.80) | 11.75 (1.83) |
|  | Median (IQR) | 6.60 (3.10) | 8.50 (3.20) | 11.90 (3.20) |
|  | Range | 2.0-9.7 | 3.4-11.9 | 6.6-16.2 |
| **Sex** |  |  |  |  |
|  | Female (%) | 1510 (50.07) | 1472 (50.12) | 1331 (50.11) |
|  | Male (%) | 1506 (49.93) | 1465 (49.88) | 1325 (49.89) |
| **Parental education** |  |  |  |  |
|  | Low (%) | 180 (5.97) | 166 (5.65) | 156 (5.87) |
|  | Medium (%) | 1337 (44.33) | 1204 (40.99) | 1172 (44.13) |
|  | High (%) | 1463 (48.51) | 1476 (50.26) | 1310 (49.32) |
| **European region of residence** |  |  |  |  |
|  | Central (%) | 1250 (41.45) | 1218 (41.47) | 1114 (41.94) |
|  | North (%) | 743 (24.64) | 721 (24.55) | 682 (25.68) |
|  | South (%) | 1023 (33.92) | 998 (33.98) | 860 (32.38) |
| **Fruit and vegetable score (%)** |  |  |  |  |
|  | Mean (SD) | 1.47 (0.75) | 1.54 (0.80) | 1.47 (0.78) |
|  | Median (IQR) | 1.38 (0.96) | 1.47 (1.02) | 1.37 (0.97) |
|  | Range | 0.00-5.71 | 0.00-5.83 | 0.00-6.07 |
|  | Missing | 58 | 154 | 106 |
| **Fiber intake (mg/kcal)** |  |  |  |  |
|  | Mean (SD) | 8.17 (1.31) | 8.23 (0.90) | 8.22 (1.27) |
|  | Median (IQR) | 8.13 (1.79) | 8.13 (1.48) | 8.07 (1.61) |
|  | Range | 3.87-15.76 | 5.76-11.56 | 4.74-13.89 |
|  | Missing | 826 | 1100 | 660 |
| **MVPA (hours/day)** |  |  |  |  |
|  | Mean (SD) | 0.67 (0.36) | 0.67 (0.36) | 0.64 (0.37) |
|  | Median (IQR) | 0.61 (0.46) | 0.62 (0.47) | 0.57 (0.47) |
|  | Range | 0.02-2.29 | 0.03-2.74 | 0.00-2.42 |
|  | Missing | 1240 | 1297 | 871 |
| **Screen time (hours/day)** |  |  |  |  |
|  | Mean (SD) | 1.60 (1.00) | 1.89 (1.08) | 2.34 (1.50) |
|  | Median (IQR) | 1.50 (1.07) | 1.75 (1.43) | 2.02 (1.79) |
|  | Range | 0.00-8.00 | 0.00-8.00 | 0.00-8.00 |
|  | Missing | 93 | 132 | 150 |
| **BMI z-scores** |  |  |  |  |
|  | Mean (SD) | 0.34 (1.16) | 0.41 (1.18) | 0.51 (1.12) |
|  | Median (IQR) | 0.23 (1.48) | 0.32 (1.67) | 0.45 (1.62) |
|  | Range | -5.42-5.80 | -5.76-4.65 | -2.96-3.83 |
|  | Obese | 204 | 214 | 179 |
| **Waist circumference z-scores** |  |  |  |  |
|  | Mean (SD) | 0.24 (1.45) | 0.59 (1.29) | 0.78 (1.25) |
|  | Median (IQR) | 0.16 (1.61) | 0.46 (1.72) | 0.71 (1.77) |
|  | Range | -27.98-5.65 | -6.79-5.33 | -7.75-4.38 |
|  | Top quartile | 461 | 443 | 316 |
|  | Missing | 76 | 22 | 55 |

Z-scores for BMI and waist circumference were calculated according to [24,25]. Boys with a BMI z-score > 2.29 and girls with a BMI z-score > 2.19 were defined as obese [24,25].

We found that the PRS-Khera provided the best prediction of BMI (see Table S1 for details on the characteristics of the other PRS). PRS-Khera was strongly associated with BMI (r^2^ = 0.11, p-value = 7.9 × 10^−81^) and waist circumference (r^2^ = 0.09, 1.8 × 10^−71^) in our study population (Table 2). Being in the top decile of the distribution of PRS-Khera was associated with 3.63 times higher odds for obesity (95% CI: [2.57, 5.14]) and with 3.09 (95% CI: [2.37, 4.03]) higher odds for being in the top quartile of waist circumference.

**Table 2.** Associations of PRS-Khera with BMI, obesity and waist circumference in IDEFICS/I.Family.

| **A) BMI** |  |  |  |  |  |  |
| --- | --- | --- | --- | --- | --- | --- |
| **Scale of PRS** | **BMI** |  |  | **Obesity** |  |  |
|  | **Est., 95% CI** | **p-value** | **R<sup>2</sup>** | **OR, 95% CI** | **p-value** | **AUC** |
| **Continuous** | 0.33 [0.30, 0.37] | 7.9e-81 | 0.108 | 2.33 [2.01, 2.70] | 2.0e-29 | 0.736 |
| **Top decile** | 0.61 [0.49, 0.73] | 5.4e-24 | 0.036 | 3.63 [2.57, 5.14] | 2.7e-13 | 0.598 |

| **B) Waist circumference** |  |  |  |  |  |  |
| --- | --- | --- | --- | --- | --- | --- |
| **Scale of PRS** | **Waist circumference** |  |  | **Waist top quartile** |  |  |
|  | **Est., 95% CI** | **p-value** | **R<sup>2</sup>** | **OR, 95% CI** | **p-value** | **AUC** |
| **Continuous** | 0.36 [0.32, 0.40] | 1.8e-71 | 0.088 | 1.97 [1.78,2.17] | 1.5e-40 | 0.683 |
| **Top decile** | 0.69 [0.55, 0.82] | 8.8e-24 | 0.032 | 3.09 [2.37,4.03] | 6.1e-17 | 0.569 |

Associations adjusted for region of residence, sex, age, parental education, vegetable score. Z-scores for BMI and waist circumference were calculated according to [24,25]. Boys with a BMI z-score > 2.29 and girls with a BMI z-score > 2.19 were defined as obese [24,25].

The correlation between PRS-Khera and BMI increased along the age range, from a squared correlation with BMI of r^2^ = 0.02 [0.01, 0.12] in the 2-year olds to r^2^ = 0.18 [0.11, 0.27] in the 14-year olds (Figure 1 and Table S2). Similar trends were found for waist circumference, for which the squared correlation with PRS-Khera was r^2^ = 0.03 [0.01, 0.07] in 3-year olds and r^2^ = 0.14 [0.08, 0.22] in 14-year olds (Figure 1 and Table S2). This increase of correlation by age group was confirmed in our sensitivity analyses using other genome-wide PRS (Figure S4 and Table S3).

**Figure 1.**
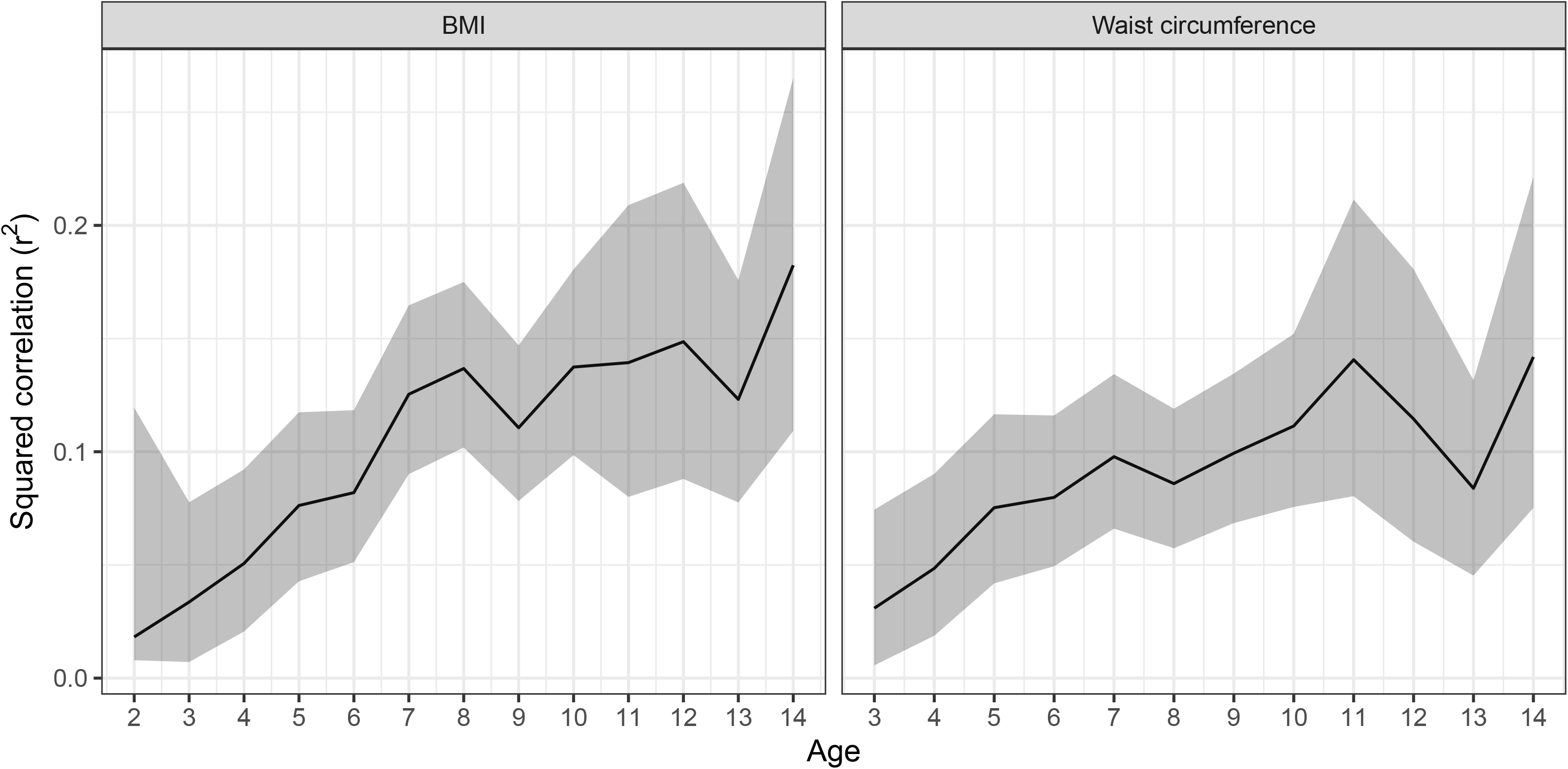
Squared correlation (r^2^ with 95% confidence intervals) of PRS-Khera with BMI and waist circumference in dependence of age. Squared correlations could not be calculated for ≥15-year old children due to the small sample size in these age groups (see Tables S1 & S2). Waist circumference was not measured in 2-year old children.

We found a significant gene-environment interaction of PRS-Khera with parental education (low vs. high) as well as with the European region of residence (Central vs. Southern) for BMI as well as for waist circumference (Figure 2, Tables S4). Children and adolescents from families with a low level of education were at a higher risk of becoming obese among those with higher genetic susceptibility than children from families with a high level of education (low: beta estimate from education-stratified analysis for association between PRS-Khera and BMI = 0.48; 95% CI: [0.38, 0.59], high: beta estimate = 0.30; 95% CI: [0.26, 0.34], adjusted p-value interaction = 0.0106, Figure 2 and Table S4). Furthermore, children and adolescents from Southern European countries showed an increased genetic susceptibility to a high BMI in comparison to children and adolescents from Central Europe (Central Europeans: beta estimate from region-stratified analysis for association between PRS-Khera and BMI = 0.29; 95% CI: [0.23, 0.34], Southern Europeans: beta estimate = 0.40; 95% CI: [0.34, 0.45], adjusted p-value interaction = 0.0246, Figure 2 and Table S4). Interactions were confirmed in our sensitivity analyses using other genome-wide PRS (Figure S5). We did not find significant interactions between PRS-Khera and sex, the comparison of low vs. medium parental education, nor the comparison of Central vs. Northern European region of residence (Figure 2, Table S4).

**Figure 2.**
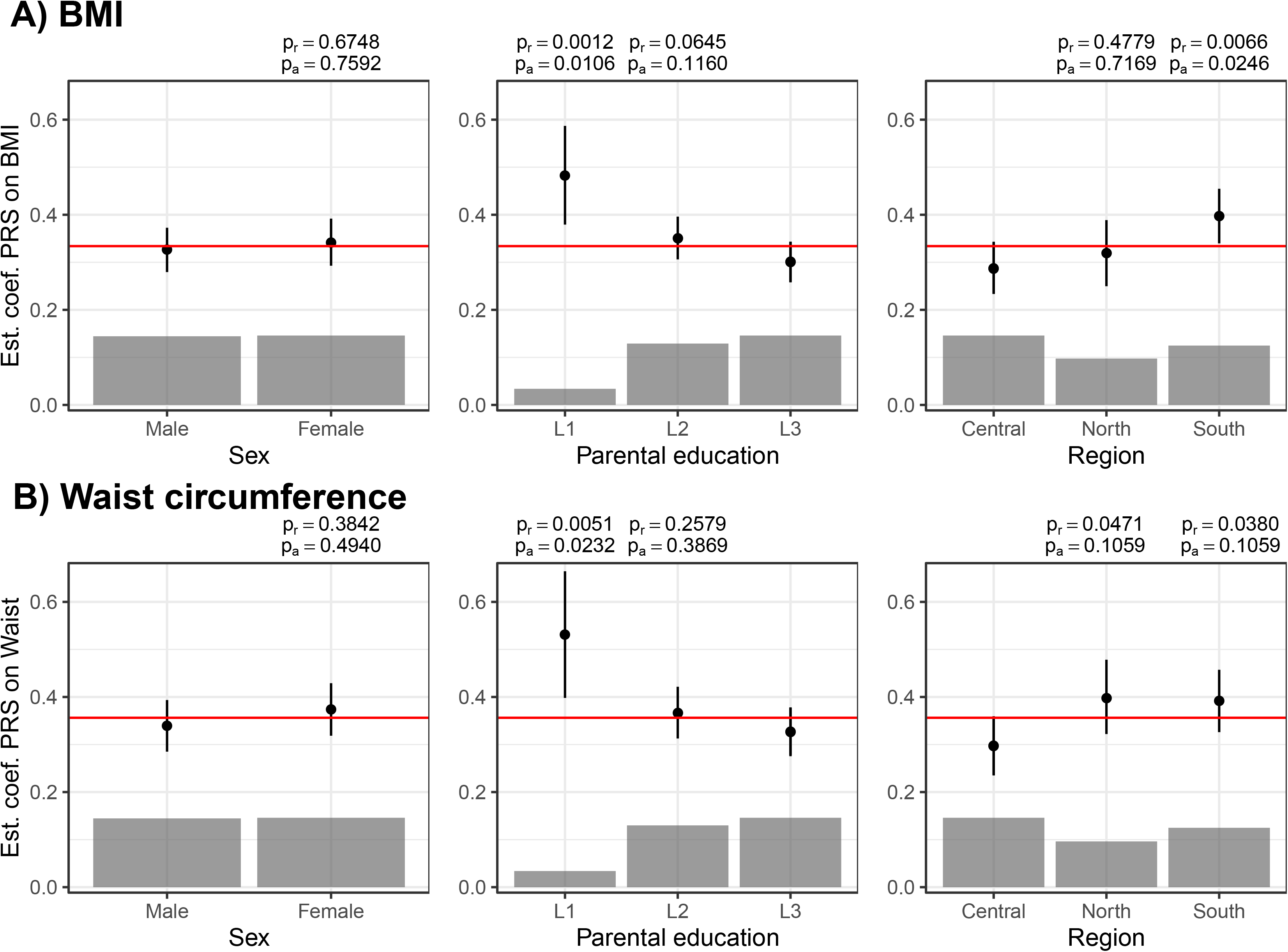
Interactions between PRS-Khera and sociodemographic factors on BMI and waist circumference. Associations between PRS and BMI / waist circumference are shown in different strata (beta estimates and 95% CIs) as well as in the whole study population (red line). Raw p-values (p_r_) and FDR-adjusted p-values (p_a_) are given for the test of deviations of the association between PRS and obesity in one subgroup in comparison to the reference category (interaction). The category without p-values is the reference category.

The genetic susceptibility to a high BMI was further modified by intake of dietary fiber and screen time (Figure 3, Tables S4). Children and adolescents with a higher fiber intake showed an attenuated risk of becoming obese despite their genetic susceptibility (adjusted p-values for interaction: 0.025 for BMI and 0.023 for waist circumference). Furthermore, the more time the children and adolescents spent in front of screens, the higher was their risk of becoming obese among those with higher genetic susceptibility (adjusted p-value interaction = 0.042). Interactions between PRS-Khera and the fruit and vegetable score or MVPA were not significant.

**Figure 3.**
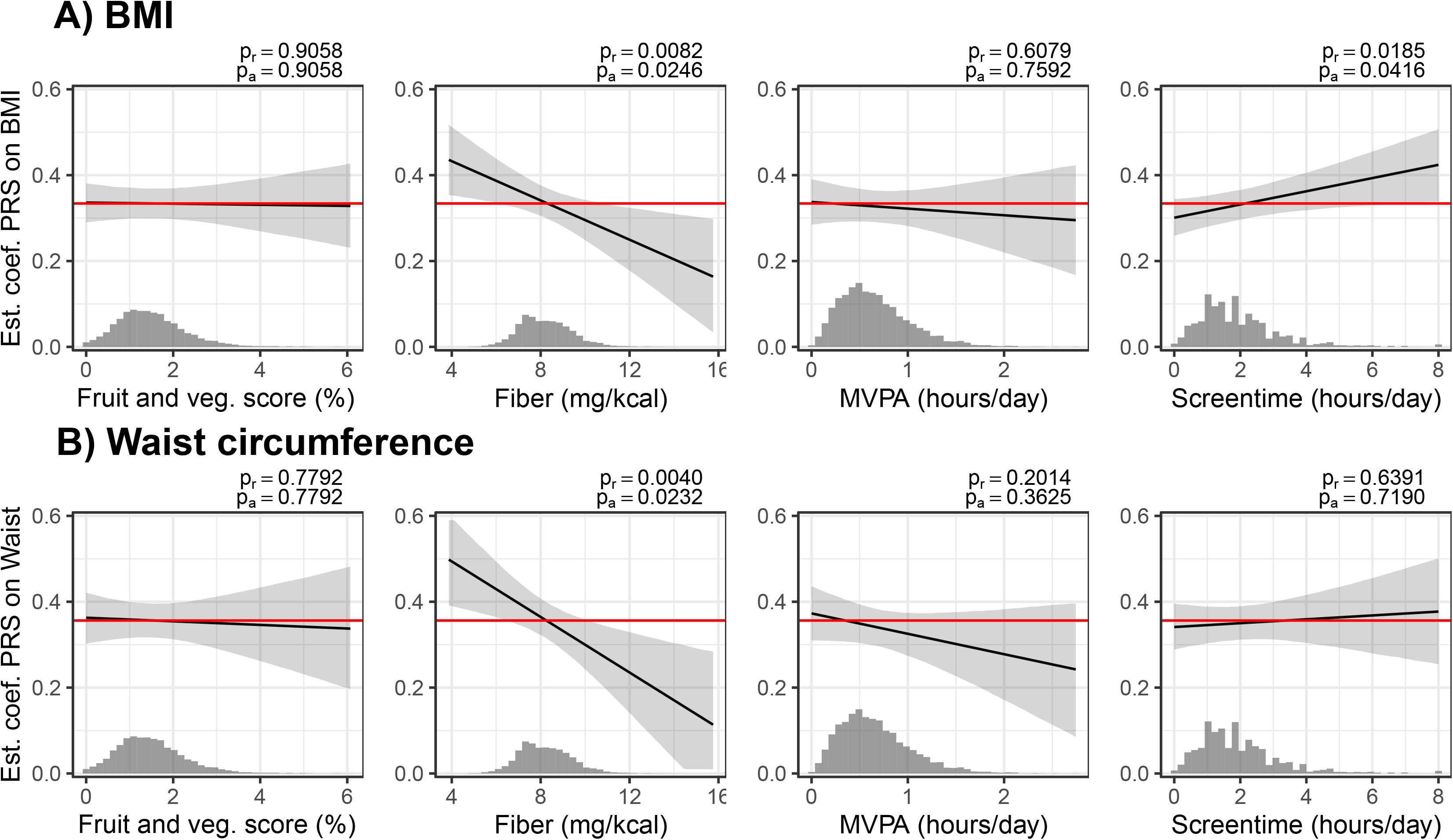
Interactions between PRS-Khera and lifestyle factors on BMI and waist circumference. Associations between PRS and obesity are shown in dependence of the PRS (beta estimates and 95% CIs) as well as in the whole study population (red line). The distributions of the lifestyle factors are shown in histograms. Raw p-values (p_r_) and FDR-adjusted p-values (p_a_) are given for the interaction terms.

## Discussion

In our pan-European cohort of children aged 2 to 16 years, we found a strong association of a polygenic risk score of obesity with BMI as well as with waist circumference and this association increased by age. We observed a prediction r^2^ of 18% in 14-year olds, which is even higher than in the original study containing mainly adults [4]. We further found significant interactions with socioeconomic and behavioral factors for BMI as well as waist circumference: we observed gene-environment interactions with (1) the European region of residence, which most likely reflect cultural lifestyle differences, (2) education, (3) dietary fiber intake and (4) the time children spent in front of screens. Of note, all of these interactions would have remained undetected in this sample of children when only focusing on genome-wide significant variants as was done in previous studies (compare Figures S5 and S6) [8–15].

### Comparison with Previous Studies

Although obesity is known to be highly polygenic, most previous gene-environment interaction analyses focused on <100 genome-wide significant variants that account for <3% of BMI variation. In this study we used a genome-wide PRS proposed in [17], which provides a more comprehensive measurement of the inherited susceptibility to obesity. Using this PRS (called PRS-Khera), we observed a prediction r^2^ of 10.8% for BMI, which is almost 5 times higher than the prediction accuracy obtained using the <100 genome-wide significant SNPs from the ~300,000 samples in [3] and twice the prediction accuracy obtained using the <1,000 genome-wide significant SNPs from the ~700,000 samples in [4] (Table S1). PRS-Khera reached a similar prediction accuracy for BMI than it has been reported from large-scale PRS in previous studies (~10.2% using the summary statistics from the ~700,000 samples and a p-value threshold of 10^−3^ (6,781 SNPs) [4] and ~8.5% [17] using a genome-wide PRS from the ~300,000 samples in [3]).

Of note, in our study, the prediction accuracy of the PRS strongly depended on age, reaching a prediction r^2^ of 18% in 14-year olds, which is in accordance with Khera et al. who showed that the association between the PRS and weight emerges early in life and increases into adulthood [17]. This surprisingly high prediction accuracy in adolescents from our study might be explained by the age difference between our study and the GIANT Consortium / UK Biobank, which was used in [4]. The GIANT Consortium / UK Biobank included mainly adults, whereas we analyzed data from children aged 2 to 16 years. In contrast to the positive correlation between age and prediction accuracy during childhood shown in this manuscript as well as in previous studies [17,46], a weak negative correlation could be observed in adults >45 years of age from the UK Biobank, an age group in which aging-related diseases become more prevalent (Table S3 in [17]). Therefore, we hypothesize that the highest prediction accuracy of the PRS for BMI might be reached in adolescents and young adults.

In our study, we found significant interactions between PRS-Khera and sociodemographic as well as lifestyle factors for BMI and waist circumference. Interactions with socioeconomic status [9], physical activity [8,9], and dietary factors [9–15] have been reported previously. However, all of these studies included only <100 genome-wide significant SNPs (e.g. from [3]). By using a genome-wide PRS we were able to detect interactions with sociodemographic and with lifestyle factors which would have remained undetected when using only genome-wide significant SNPs (Figures S5 and S6).

Furthermore, previous GxE interaction studies [8–15] were mainly based on adults whereas in our study we analyzed data from children aged 2 to 16 years. Therefore, our results provide new insights about how a healthy childhood environment might partly offset a genetic predisposition to obesity during childhood and adolescence. In our study, we identified children from families with low levels of education as being about 61% more susceptible to the polygenic burden of obesity than children from families with a high level of education. In addition, we found that children from Southern Europe had a higher genetic susceptibility to obesity in comparison to children from Central Europe. Parental education and region of residence reflect a variety of social and cultural differences and many of them are difficult to capture by questionnaires. Since a previous analysis of the same cohort showed that low parental education was associated with higher intakes of unhealthy food among children, e.g. sugar-rich and fatty foods [47,48], part of the effect modification might be due to dietary habits. The differences in the risk of becoming obese among children with a higher genetic susceptibility across different European regions might be explained by differences in dietary or cultural habits [49,50].

Furthermore, we found an interaction between PRS-Khera and intake of fiber, with children with a higher intake of fiber having a reduced risk for obesity despite their genetic susceptibility. This finding is in line with many other studies that have shown that a healthy diet can attenuate the genetic burden of obesity [9–15]. Interactions between PRS-Khera and physical activity (MVPA) were not significant, but the direction of interaction effect was in line with previous studies [8,9]. An explanation for this might be that MVPA was only assessed in ~40% of our analysis group (Table 1), which reduced the statistical power to detect interactions between MVPA and PRS.

### Strengths and Limitations of this Study

Important strengths of this study include: detailed and repeated phenotyping of participants in this cohort with partly objective measures (MVPA), inclusion of thousands of children from diverse regions in Europe and the longitudinal approach across key developmental periods [22]. Dietary assessment in children is a challenging task [51], and different dietary assessment have different strengths and limitations. We used two different dietary assessment methods – a fruit and vegetable score derived from FFQs and fiber intake calculated from the more detailed 24-hour dietary recalls. The harmonized protocol in all countries that was enforced by a central quality control and a central data management ensures comparability of measurements across study centers. Another major strength of our study is the application of genome-wide PRS for obesity, which has an almost 5 times higher prediction accuracy than previously used PRS [9–15] and with which we identified interactions that would have remained undetected when only focusing on genome-wide significant variants (compare Figures S5 and S6).

Our study also has several limitations. First, measurement errors of self-reported lifestyle behaviors are inevitable. However, measurement error in environmental exposure typically biases the interaction effect toward the null [52], which does not increase the risk for false positive findings but reduces the statistical power to detect subtle interactions. Second, the use of PRS derived from associations with BMI in the analyses of waist circumference led to slightly lower prediction accuracy for waist circumference than for BMI. However, since PRS-Khera is known to be a strong risk factor for severe obesity and associated health outcomes [17], we decided to use this PRS for both obesity measurements.

### Conclusions

Our study showed significant interactions between the polygenic risk for an increased BMI and sociodemographic and behavioral factors that affect BMI as well as waist circumference. Among children with a high genetic risk, we identified children from Southern Europe, children from families with a low level of education, children with a low intake of fiber and children who spend more time in front of screens as being particularly susceptible to obesity. These results provide evidence that the risk for obesity among children with a high genetic susceptibility varies by environmental and sociodemographic factors during childhood. This has important implications for future public health prevention efforts, because it suggests that children at a high genetic risk may benefit even more from prevention measures than children with a low genetic risk.

## Acknowledgments

The authors wish to thank the IDEFICS children and their parents for participating in this extensive examination. We are grateful for the support of school boards, head teachers and communities, and for the effort of the study nurses, interviewers, data managers and, laboratory technicians.

