## Supplementary Material for "A healthy childhood environment helps to combat inherited susceptibility to obesity"

### Methods

#### *Genotyping and Quality Control*

DNA was extracted from saliva or blood samples using established procedures. Genotyping of 3,515 children was performed on the UK Biobank Axiom array (Santa Clara, USA) in two batches (2015 and 2017). Following the recommendations of [1], sample and genotype quality control measures were applied. After visual inspection of the cumulative SNP coverage distribution, SNPs with call rates less than 97.5%, relative minor allele frequency (MAF) < 0.05 (batch1) and 0.08 (batch2), not being in Hardy-Weinberg equilibrium (p-value < $10^{-4}$.) were excluded. Children with call rates less than 98% (batch1) and 96% (batch2), sex mismatch, anomalously high heterozygosity [1], duplicates, or population outliers were also excluded from the sample. We performed principal components analysis (PCA) to identify population outliers using the R package SNPRelate [2]. A sample of 3,099 children and 570,587 (batch1) / 551,909 (batch2) SNPs, respectively, remained for further analyses. Genome-wide imputation was conducted using Minimac3 (https://genome.sph.umich.edu/wiki/Minimac3) and reference haplotypes from unrelated individuals from the 1000 Genomes Project phase III v5. A total of 3,424,677 imputed genotypes with an estimated posterior genotype probability > 0.8 and MAF ≥ 0.05 from 3,099 children and adolescents were used in the analyses.

#### *Polygenic Risk Score Calculation*

In our sensitivity analyses, we calculated four PRS for BMI - two PRS based on genome-wide summary statistics of European ancestry populations and two based on only genome-wide significant SNPs from the reference populations.

Two PRS were calculated using PRSice [3], one using the same reference population as for PRS-Khera (PRS-Locke$^{BT}$; BT: best threshold) and a second one based on summary statistics from the largest published GWAS study of BMI to date (~700,000 samples) [4] (PRS-Yengo$^{BT}$). PRSice calculates the sum of alleles weighted by their effect sizes estimated from a GWAS of that phenotype in an independent sample (here from [4]) using an approach called clumping and thresholding. Clumping was used to obtain SNPs in linkage equilibrium with an $r^2 < 0.1$ within a 500 kbp window, keeping the SNP with the lower p-value observed in the external dataset for the analysis. Multiple scores were then created according to the significance of their association with the phenotype. As proposed in [3], the PRS that predicted the phenotype (BMI) best (highest $R^2$) was used for analysis, resulting in a p-value threshold of 1 for PRS-Locke$^{BT}$ (Figure S1) and 0.5 for PRS-Yengo$^{BT}$ (Figure S2).

The two PRS that were based on only genome-wide significant (GS) SNPs from the reference populations (PRS-Locke$^{GS}$ with significant SNPs from [5] and PRS-Yengo$^{GS}$ with significant SNPs from [4]) were also calculated using PRSice by setting a fixed p-value threshold of $5 \times 10^{-8}$.

*Assessment of Dietary Intake*

Information on dietary intake was collected by two retrospective methods, repeated 24-hour dietary recalls (24HDRs) and food frequency questionnaires (FFQs). For the present analysis, fruit and vegetable intake was obtained from the FFQs, which were self-reported for adolescents 12 years and older and proxy-reported by a parent or other caregiver for children below the age of 12 years. In the IDEFICS study, the FFQ asked for the usual consumption frequency of 43 food and beverage items, out of which 4 were fruit and vegetable items including cooked vegetables, potatoes and beans; raw vegetables (mixed salad, carrot, fennel, cucumber, lettuce, tomato, and local examples); fresh fruits (also freshly squeezed, fruit smoothie) without added sugar; and fresh fruits (also freshly squeezed, fruit smoothie) with added sugar. In the I.Family study, the FFQ was extended to include 59 items for children and 60 item for adolescents, out of which 5 were fruit and vegetables items including potatoes (cooked, not fried); other cooked vegetables (with local examples); raw vegetables (mixed salad, carrot, fennel, cucumber, lettuce, tomato, and local examples); fresh fruits (also as freshly squeezed juice) without added sugar; and fresh fruits (also as freshly squeezed juice) with added sugar. The FFQs in the IDEFICS/I.Family studies asked for the usual consumption frequency by the following question: "In the last month, how many times did you eat or drink the following food items?" Possible response categories were recoded to weekly consumption frequencies as follows: never/less than once a week (recoded to 0), 1–3 times a week (recoded to 2), 4–6 times a week (recoded to 5), 1 time per day (recoded to 7), 2 times per day (recoded to 14), 3 times per day (recoded to 21) or 4 or more times per day (recoded to 30). To maintain comparability across countries, the same foods and beverages were translated into eight languages, but for some food items, country-specific foods were also noted as examples. The items from the FFQ have been found to be reproducible with a weighted kappa coefficients of 0.49 and a Spearman's correlation coefficient of 0.59 for vegetables [6], and a validation study of the FFQ against repeated 24HDRs showed that the gross misclassification for fruit and vegetables was below 5% [7]. We expressed fruit and vegetable consumption as relative frequency of all consumed foods in the FFQ [8]. Items of the FFQ that were omitted were assumed as not consumed if less than half of the items on the FFQ were missing; otherwise, the consumption frequency was set to missing [8,9].

Energy and dietary fiber intake were assessed by using repeated 24HDRs [10,11]. The offline software called SACINA (Self-Administered Children and Infant Nutrition Assessment) used in the IDEFICS study was amended and delivered as a web-based version, called SACANA (Self-Administered Child, Adolescent and Adult Nutrition Assessment) in the I.Family study. In the IDEFICS study, parents or other caregivers proxy-reported 24HDRs for their children. In I.Family, children from 8 years of age onwards self-completed their 24HDRs, while parental assistance was given at younger ages. Participants were assisted by trained survey personnel or dieticians when completing the first 24HDR and asked to recall and enter all foods and beverages (types and amounts) consumed during the previous day, starting with the first meal in the morning after waking up. Standardized photographs were displayed on the screen to assist portion size estimation. Food items from SACINA and SACANA were linked with country-specific food composition tables to calculate energy and nutrient intakes. Usual intakes for fiber were estimated based on the validated National Cancer Institute (NCI) method, which is one of the most widely accepted methods for this purpose [12,13]. This method allows for the inclusion of covariates such as age and accounts for different intakes on weekend days vs. weekdays, and further corrects for the day-to-day variation in energy and fiber intakes. Usual intakes were estimated for each child stratified by sex and considering age as a covariate. Fiber intake was here expressed in relation to total energy intake in mg/kcal.

*Assessment of Physical Activity*

Physical activity was objectively measured by using Actigraph's uniaxial or three-axial accelerometers [14]. At baseline and FU1, children were asked to wear the accelerometer for 3 days (including one weekend day) and at FU2 for a full week during waking hours (except when swimming or showering), so that a full day of physical activity could be assessed. The accelerometers were attached to the right hip with an elastic belt. Participants (either the parents or the adolescents themselves) were given written instructions on how to use the accelerometer and were asked to complete diaries to record non-wear times of the device. The daily average cumulative duration of time spent performing moderate-to-vigorous physical activity (MVPA) was expressed as minutes per day according to the cut-off value by Evenson et al. [15]. Time spent in MVPA is based on cleaned accelerometer data that only contain measurements that have passed the minimum wear time criteria of at least 3 measurement days and at least 360 minutes of valid time per day. Although it does not cover all waking hours, a monitoring period of 360 minutes or more has been shown to provide an acceptable reliability in young children [16,17]. The valid wearing time of the included sample was longer in practice (mean 12.0 (standard deviation (SD)

1.6) h/d). All accelerometer recordings were integrated over 60 s epochs. Non-wear time was defined as 20 min or more of consecutive 0 counts. The average activity level of the children was defined by counts per minute (cpm). The average activity level of the children was defined by counts per minute per day (cpm/d, average over all valid days).

*Assessment of Screen Time*

As a proxy indicator for sedentary time, screen time duration was assessed by asking how many hours per day the child/adolescent usually spends watching television (including videos or DVDs) and by another question on the time sitting in front of a computer and game console [18,19]. The first question was included in a pilot study testing the reproducibility of the questionnaire, and the results suggested agreement between repeat measures (Cohen's kappa: k = 0.85 (95% CI: [0.78,0.91]) [20]. For both questions, separate answers were given for weekdays and weekend days. The answer categories 'Not at all', 'less than 30 min. per day', '30 min. to 1 hour per day', 'about 1-2 hours per day', 'about 2-3 hours per day' and 'more than 3 hours per day' were recoded to daily frequencies with the conversion factors 0, 0.25, 0.75, 1.5, 2.5 and 4, respectively. Responses were weighted and summed over weekdays and weekend days and the quantified frequencies from both questions were added to create a continuous variable of total screen time in hours per day. Parents proxy-reported for their children at baseline and FU1, and for children <12 at FU2, while children ≥ 12 at FU2 self-reported their screen time.

**Comparison of PRS-Khera with other PRS for BMI**

The three PRS that were based on genome-wide summary statistics (Khera, $Locke^{BT}$ and $Yengo^{BT}$) were moderately correlated (Spearman correlation between 0.61 and 0.66), whereas correlations with the PRS that were based on only genome-wide significant SNPs ($Locke^{GS}$ and $Yengo^{GS}$) were much lower (0.16 to 0.56) (Figure S3).

We found that the PRS-Khera provided the best prediction of BMI ($r^2$=0.11, p-value = 7.9 x $10^{-81}$) (Table 2). These associations were very robust across different reference populations (300,000 samples from [5] versus 700,000 samples from [4]) and across different genome-wide PRS approaches (PRSice vs. LDPred) (Table S1). In contrast, previous PRS for BMI, that were only based on the 77 genome-wide significant SNPs from [5], explained a much smaller part of the variance of BMI (PRS-$Locke^{GS}$: $r^2$=0.02, p-value = 4.2 x $10^{-29}$). Furthermore, the association strongly depended on the sample size of the reference population. Using the genome-wide significant SNPs from the 700,000 samples from [4] instead of the 300,000 samples from [5], increased the prediction accuracy of both BMI and waist circumference from $r^2$ = 0.02 to 0.06 for BMI and from $r^2$ = 0.02 to 0.05 for waist circumference.

The increase in correlations between PRS and BMI by age was very robust across different genome-wide PRS approaches (Khera, $Locke^{BT}$, $Yengo^{BT}$), but less pronounced for the PRS that were only based on genome-wide significant SNPs ($Locke^{GS}$ and $Yengo^{GS}$) (Figure S4 and Table S3). The squared correlation of $Locke^{GS}$ with BMI was about the same for every age group from $r^2$ = 0.01 [0.00, 0.03] to $r^2$ = 0.03 [0.00, 0.07] and the squared correlation of $Yengo^{GS}$ with BMI increased from $r^2$ = 0.05 [0.02, 0.11] to $r^2$ = 0.11 [0.04, 0.17].

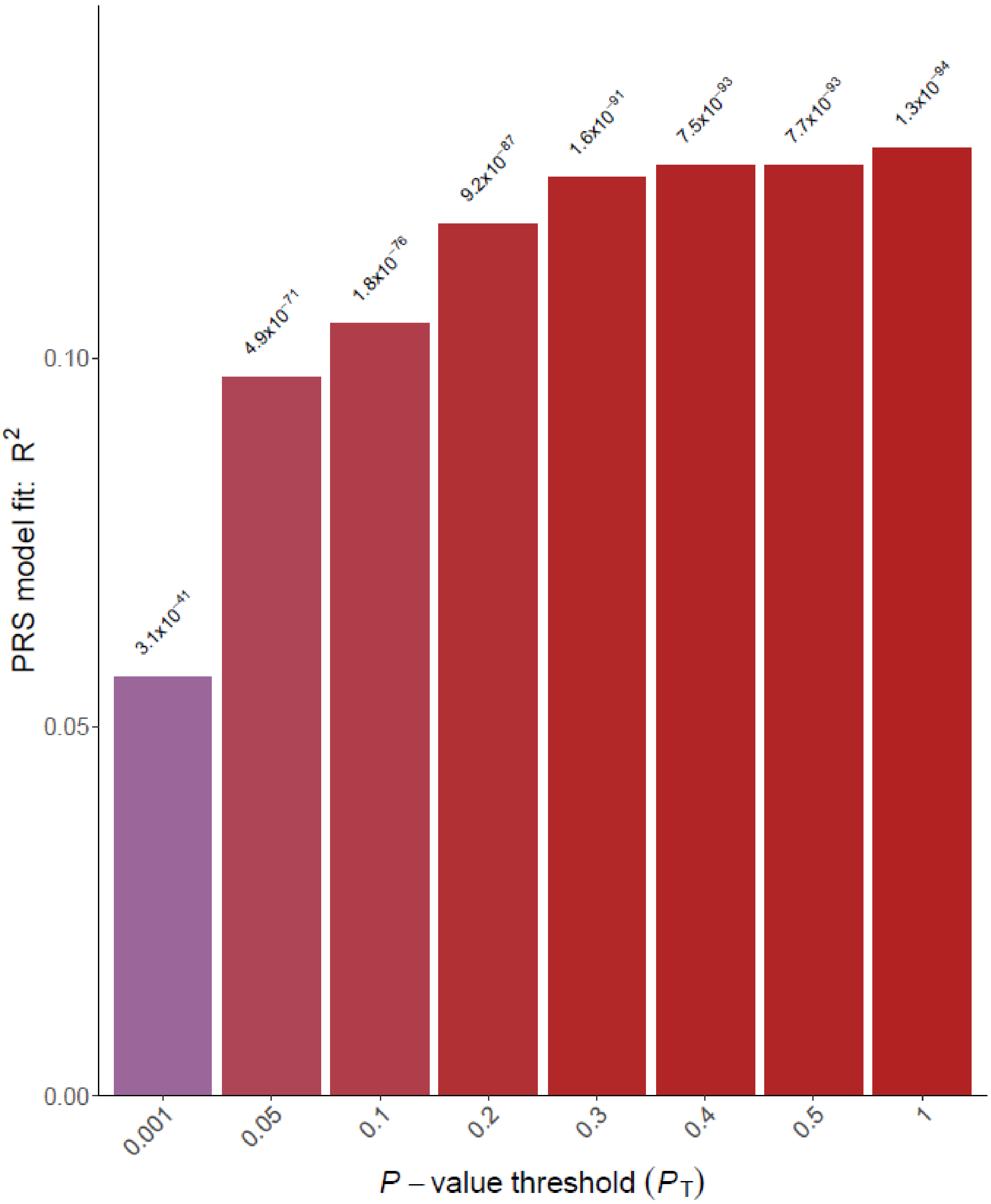


**Figure S1.** Association of PRS-Locke with BMI in dependence of the p-value threshold chosen for the inclusion of SNPs. The p-values for the associations between the different PRS and BMI are stated above the bars. The highest prediction capacity ($R^2$) was reached when all 80,975 independent SNPs were included in the PRS (p-value threshold of 1). This PRS (PRS-Locke$^{BT}$; BT: best threshold) was used for all subsequent analyses.

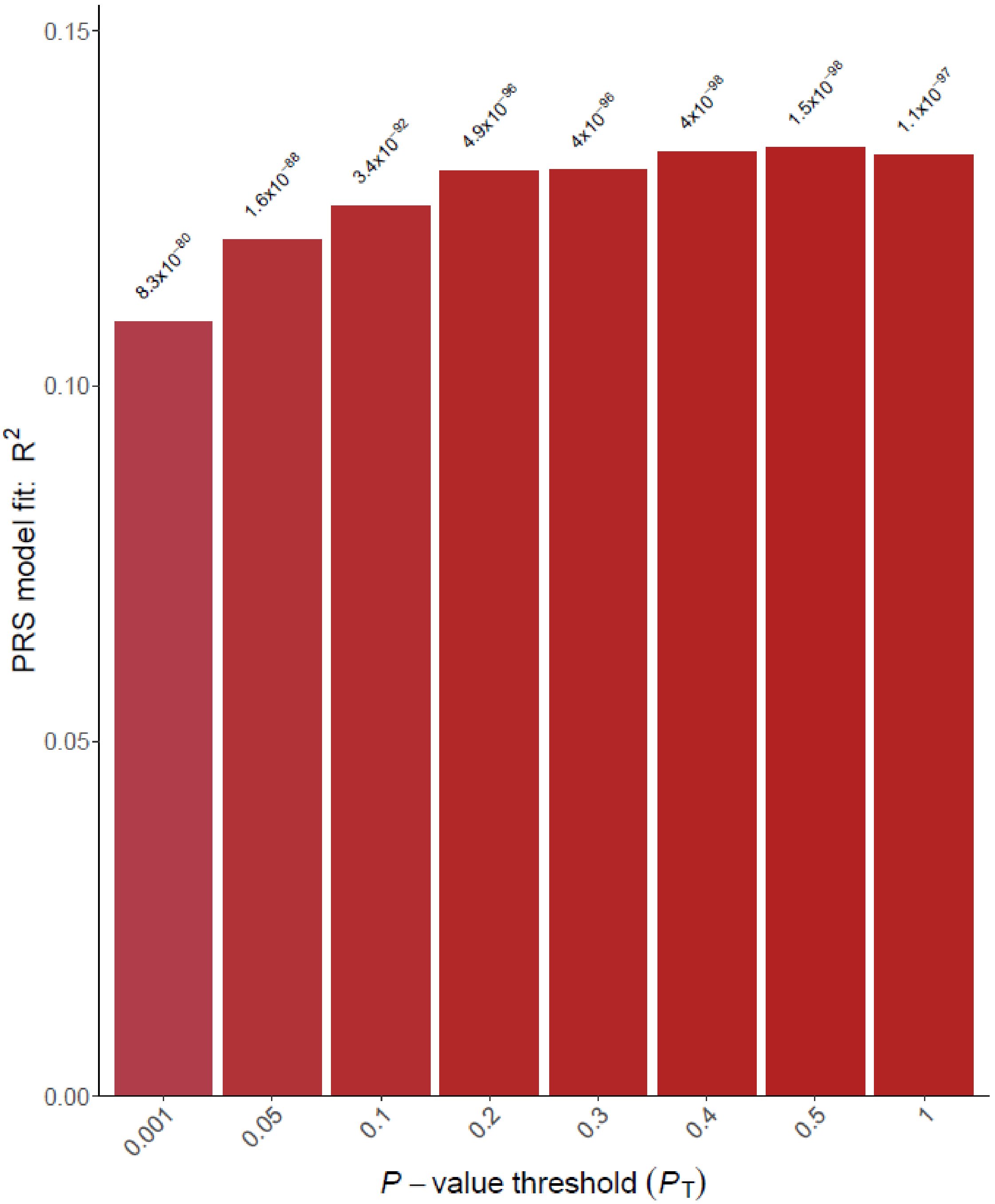


**Figure S2.** Association of PRS-Yengo with BMI in dependence of the p-value threshold chosen for the inclusion of SNPs. The p-values for the associations between the different PRS and BMI are stated above the bars. The highest prediction capacity ($R^2$) was reached for a p-value threshold of 0.5. This PRS (PRS-Yengo$^{BT}$; BT: best threshold) includes 63,523 independent SNPs and was used for all subsequent analyses.

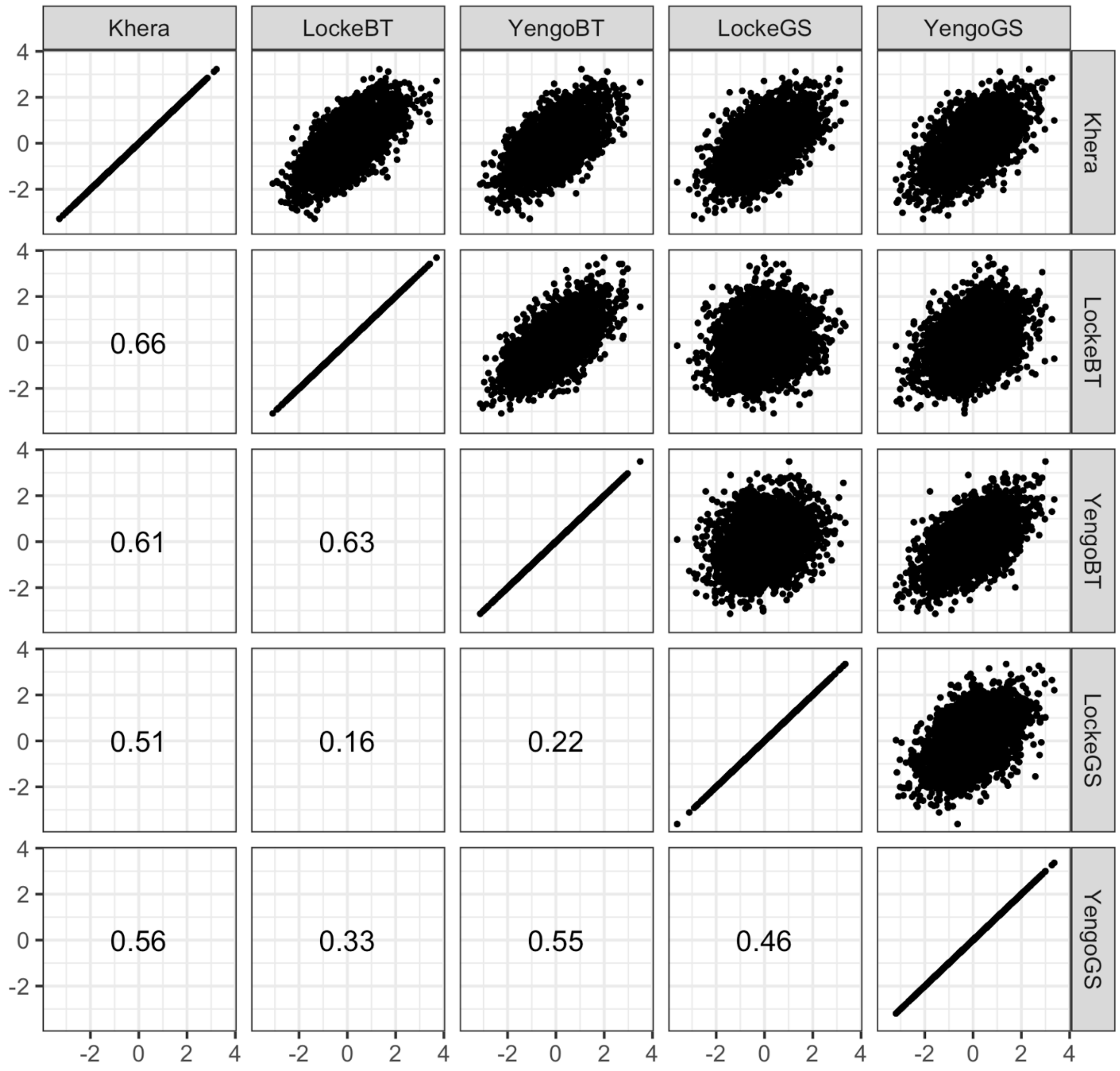


**Figure S3.** Spearman correlations between different PRS. The scatter plots show the pairwise comparisons of different PRS for each of the 3,098 study participants.

**Table S1. Associations of PRS with BMI and obesity in IDEFICS/I.Family.**

| | | | BMI | | | Obesity | | |
|---|---|---|---|---|---|---|---|---|
| **Scale of PRS** | **Ref. pop. (N)** | **PRS** | **Est., 95% CI** | **p-value** | **$R^2$** | **OR, 95% CI** | **p-value** | **AUC** |
| **Continuous** | Locke (~300,000) | Khera | 0.33 [ 0.30, 0.37] | 7.9e-81 | 0.108 | 2.33 [2.01, 2.70] | 2.0e-29 | 0.736 |
| | Locke (~300,000) | $Locke^{BT}$ | 0.29 [ 0.25, 0.33] | 6.7e-55 | 0.102 | 2.17 [1.87, 2.53] | 8.9e-24 | 0.750 |
| | Yengo (~700,000) | $Yengo^{BT}$ | 0.34 [ 0.30, 0.37] | 3.7e-81 | 0.107 | 2.23 [1.93, 2.59] | 2.0e-26 | 0.723 |
| | Locke (~300,000) | $Locke^{GS}$ | 0.20 [ 0.17, 0.24] | 4.2e-29 | 0.024 | 1.68 [1.47, 1.92] | 3.6e-14 | 0.612 |
| | Yengo (~700,000) | $Yengo^{GS}$ | 0.28 [ 0.25, 0.32] | 7.2e-58 | 0.061 | 1.87 [1.62, 2.15] | 8.1e-18 | 0.648 |
| **Top decile** | Locke (~300,000) | Khera | 0.61 [ 0.49, 0.73] | 5.4e-24 | 0.036 | 3.63 [2.57, 5.14] | 2.7e-13 | 0.598 |
| | Locke (~300,000) | $Locke^{BT}$ | 0.64 [ 0.52, 0.76] | 2.1e-25 | 0.047 | 4.54 [3.25, 6.34] | 8.9e-19 | 0.633 |
| | Yengo (~700,000) | $Yengo^{BT}$ | 0.57 [ 0.45, 0.69] | 9.8e-21 | 0.028 | 2.69 [1.90, 3.82] | 2.9e-08 | 0.564 |
| | Locke (~300,000) | $Locke^{GS}$ | 0.40 [ 0.28, 0.51] | 5.9e-11 | 0.008 | 2.67 [1.84, 3.87] | 2.2e-07 | 0.541 |
| | Yengo (~700,000) | $Yengo^{GS}$ | 0.59 [ 0.47, 0.71] | 1.6e-22 | 0.021 | 3.04 [2.12, 4.36] | 1.7e-09 | 0.554 |

Associations adjusted for region of residence, sex, age, parental education, vegetable score. Z-scores for BMI were calculated according to [21,22]. Boys with a BMI z-score > 2.29 and girls with a BMI z-score > 2.19 were defined as obese [21,22]. BT: PRSice with best p-value threshold, GS: only genome-wide significant SNPs from the reference population were included; reference populations: Locke [5], Yengo [4].

**Table S2.** Squared correlations ($r^2$) between PRS-Khera and BMI & waist circumference in dependence of age.

| | BMI | | Waist circumference | |
|---|---|---|---|---|
| **Age** | **N** | **$r^2$ [95% CI]** | **N** | **$r^2$ [95% CI]** |
| 2 | 79 | 0.02 [0.01, 0.12] | 0 | NA [NA, NA] |
| 3 | 381 | 0.03 [0.01, 0.08] | 370 | 0.03 [0.01, 0.07] |
| 4 | 541 | 0.05 [0.02, 0.09] | 521 | 0.05 [0.02, 0.09] |
| 5 | 712 | 0.08 [0.04, 0.12] | 703 | 0.08 [0.04, 0.12] |
| 6 | 937 | 0.08 [0.05, 0.12] | 931 | 0.08 [0.05, 0.12] |
| 7 | 1055 | 0.13 [0.09, 0.16] | 1047 | 0.10 [0.07, 0.13] |
| 8 | 1167 | 0.14 [0.10, 0.17] | 1156 | 0.09 [0.06, 0.12] |
| 9 | 1134 | 0.11 [0.08, 0.15] | 1130 | 0.10 [0.07, 0.13] |
| 10 | 930 | 0.14 [0.10, 0.18] | 919 | 0.11 [0.08, 0.15] |
| 11 | 377 | 0.14 [0.08, 0.21] | 367 | 0.14 [0.08, 0.21] |
| 12 | 382 | 0.15 [0.09, 0.22] | 375 | 0.11 [0.06, 0.18] |
| 13 | 599 | 0.12 [0.08, 0.18] | 576 | 0.08 [0.05, 0.13] |
| 14 | 304 | 0.18 [0.11, 0.27] | 295 | 0.14 [0.08, 0.22] |
| 15 | 10 | NA [NA, NA] | 9 | NA [NA, NA] |
| 16 | 1 | NA [NA, NA] | 1 | NA [NA, NA] |

Squared correlations could not be calculated for ≥15-year old children due to the small sample size in these age groups. Waist circumference was not measured in 2-year old children.

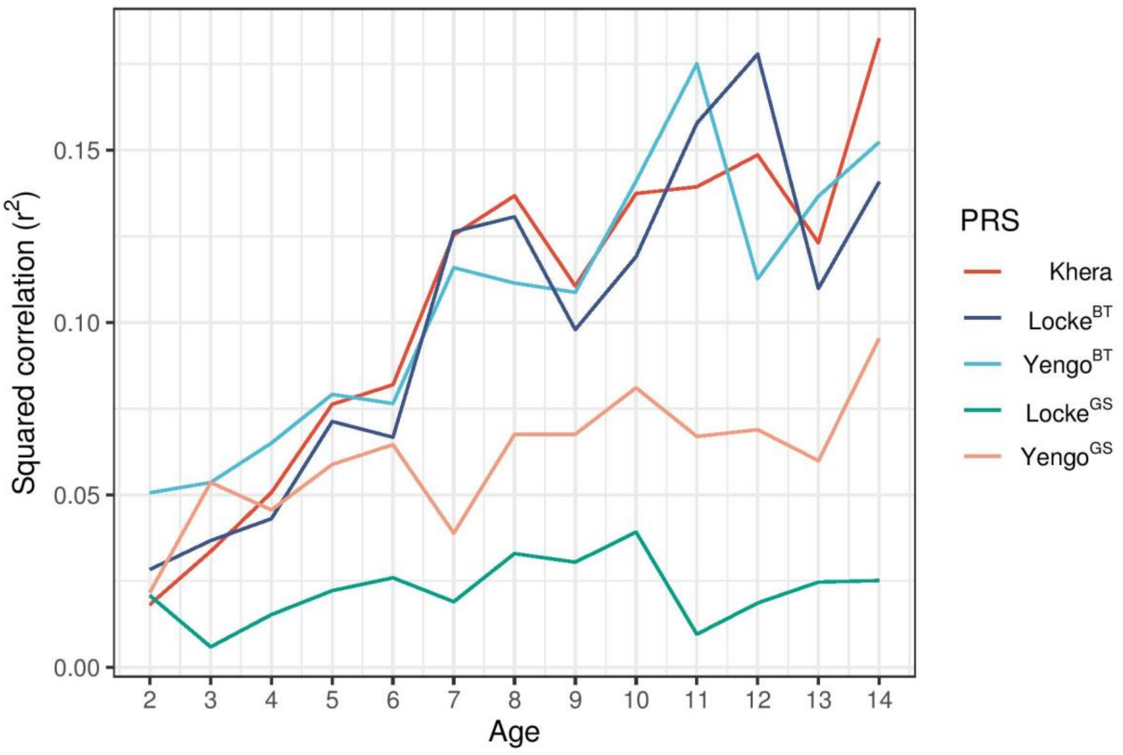


**Figure S4. Squared correlation ($r^2$) of the different PRS with BMI in dependence of age.** Squared correlations could not be calculated for ≥15-year old children due to the small sample size in these age groups (see Table S2).

**Table S3.** Squared correlations ($r^2$) [95% CI] between the different PRS and BMI in dependence of age.

| **Age** | **N** | **Khera** | **Locke$^{BT}$** | **Yengo$^{BT}$** | **Locke$^{GS}$** | **Yengo$^{GS}$** |
|---|---|---|---|---|---|---|
| 2 | 79 | 0.02 [0.01, 0.12] | 0.03 [0.00, 0.14] | 0.05 [0.00, 0.18] | 0.02 [0.01, 0.13] | 0.02 [0.01, 0.13] |
| 3 | 381 | 0.03 [0.01, 0.08] | 0.04 [0.01, 0.08] | 0.05 [0.02, 0.11] | 0.01 [0.00, 0.03] | 0.05 [0.02, 0.11] |
| 4 | 541 | 0.05 [0.02, 0.09] | 0.04 [0.02, 0.08] | 0.07 [0.03, 0.11] | 0.02 [0.00, 0.04] | 0.05 [0.02, 0.09] |
| 5 | 712 | 0.08 [0.04, 0.12] | 0.07 [0.04, 0.11] | 0.08 [0.05, 0.12] | 0.02 [0.01, 0.05] | 0.06 [0.03, 0.10] |
| 6 | 937 | 0.08 [0.05, 0.12] | 0.07 [0.04, 0.10] | 0.08 [0.05, 0.11] | 0.03 [0.01, 0.05] | 0.06 [0.04, 0.10] |
| 7 | 1055 | 0.13 [0.09, 0.16] | 0.13 [0.09, 0.17] | 0.12 [0.08, 0.15] | 0.02 [0.01, 0.04] | 0.04 [0.02, 0.06] |
| 8 | 1167 | 0.14 [0.10, 0.17] | 0.13 [0.10, 0.17] | 0.11 [0.08, 0.15] | 0.03 [0.02, 0.06] | 0.07 [0.04, 0.10] |
| 9 | 1134 | 0.11 [0.08, 0.15] | 0.10 [0.07, 0.13] | 0.11 [0.08, 0.14] | 0.03 [0.01, 0.05] | 0.07 [0.04, 0.10] |
| 10 | 930 | 0.14 [0.10, 0.18] | 0.12 [0.08, 0.16] | 0.14 [0.10, 0.18] | 0.04 [0.02, 0.07] | 0.08 [0.05, 0.12] |
| 11 | 377 | 0.14 [0.08, 0.21] | 0.16 [0.10, 0.23] | 0.18 [0.11, 0.25] | 0.01 [0.00, 0.04] | 0.07 [0.03, 0.12] |
| 12 | 382 | 0.15 [0.09, 0.22] | 0.18 [0.11, 0.25] | 0.11 [0.06, 0.18] | 0.02 [0.00, 0.05] | 0.07 [0.03, 0.12] |
| 13 | 599 | 0.12 [0.08, 0.18] | 0.11 [0.07, 0.16] | 0.14 [0.09, 0.19] | 0.02 [0.01, 0.05] | 0.06 [0.03, 0.10] |
| 14 | 304 | 0.18 [0.11, 0.27] | 0.14 [0.08, 0.22] | 0.15 [0.08, 0.23] | 0.03 [0.00, 0.07] | 0.10 [0.04, 0.17] |
| 15 | 10 | NA [NA, NA] | NA [NA, NA] | NA [NA, NA] | NA [NA, NA] | NA [NA, NA] |
| 16 | 1 | NA [NA, NA] | NA [NA, NA] | NA [NA, NA] | NA [NA, NA] | NA [NA, NA] |

Squared correlations could not be calculated for ≥15-year old children due to the small sample size in these age groups.

**Table S4. Interactions between PRS-Khera and parental education for BMI and waist circumference.**

**A) BMI**

| **Sociodemographic variable** | **Category** | **Est., 95% CI** | **Raw p-value** | **Adjusted p-value** |
|---|---|---|---|---|
| Sex | Male | 0.33 [0.28,0.37] | Reference | |
| | Female | 0.34 [0.29,0.39] | 0.6748 | 0.7592 |
| Parental education | Low | 0.48 [0.38,0.59] | **0.0012** | **0.0106** |
| | Medium | 0.35 [0.31,0.39] | 0.0645 | 0.1160 |
| | High | 0.30 [0.26,0.34] | Reference | |
| Region | Central | 0.29 [0.23,0.34] | Reference | |
| | North | 0.32 [0.25,0.39] | 0.4779 | 0.7169 |
| | South | 0.40 [0.34,0.45] | **0.0066** | **0.0246** |

**B) Waist circumference**

| **PRS** | **Category** | **Est., 95% CI** | **Raw p-value** | **Adjusted p-value** |
|---|---|---|---|---|
| Sex | Male | 0.34 [0.29,0.39] | Reference | |
| | Female | 0.37 [0.32,0.43] | 0.3842 | 0.4940 |
| Parental education | Low | 0.53 [0.40,0.67] | **0.0051** | **0.0232** |
| | Medium | 0.37 [0.31,0.42] | 0.2579 | 0.3869 |
| | High | 0.33 [0.27,0.38] | Reference | |
| Region | Central | 0.30 [0.24,0.36] | Reference | |
| | North | 0.40 [0.32,0.47] | **0.0471** | 0.1059 |
| | South | 0.39 [0.33,0.46] | **0.0380** | 0.1059 |

Associations between PRS and obesity are shown in different strata (beta estimates and 95% confidence intervals). P-values are given for the test of deviations of the association between PRS and obesity in one subgroup in comparison to the reference category (interaction). P-values were adjusted according to the number of tested environmental factors using the false-discovery rate (FDR).

**Table S5. Interaction between PRS-Khera and lifestyle factors for BMI and waist circumference.**

**A) BMI**

| Environmental factor | Est., 95% CI | Raw p-value | Adjusted p-value |
|---|---|---|---|
| Fruit and vegetable score | 0.00 [-0.02, 0.02] | 0.9058 | 0.9058 |
| Fiber intake | -0.02 [-0.04,-0.01] | **0.0082** | **0.0246** |
| MVPA | -0.01 [-0.07, 0.04] | 0.6079 | 0.7592 |
| Screen time | 0.02 [ 0.00, 0.03] | **0.0185** | **0.0416** |

**B) Waist circumference**

| Environmental factor | Est., 95% CI | Raw p-value | Adjusted p-value |
|---|---|---|---|
| Fruit and vegetable score | 0.00 [-0.03, 0.02] | 0.7792 | 0.7792 |
| Fiber intake | -0.03 [-0.06,-0.01] | **0.0040** | **0.0232** |
| MVPA | -0.05 [-0.12, 0.03] | 0.2014 | 0.3625 |
| Screen time | 0.00 [-0.01, 0.02] | 0.6391 | 0.7190 |

Beta estimates, 95% confidence intervals and p-values are given for the interaction between the PRS and each environmental factor with BMI. P-values were adjusted according to the number of tested environmental factors using the false-discovery rate (FDR).

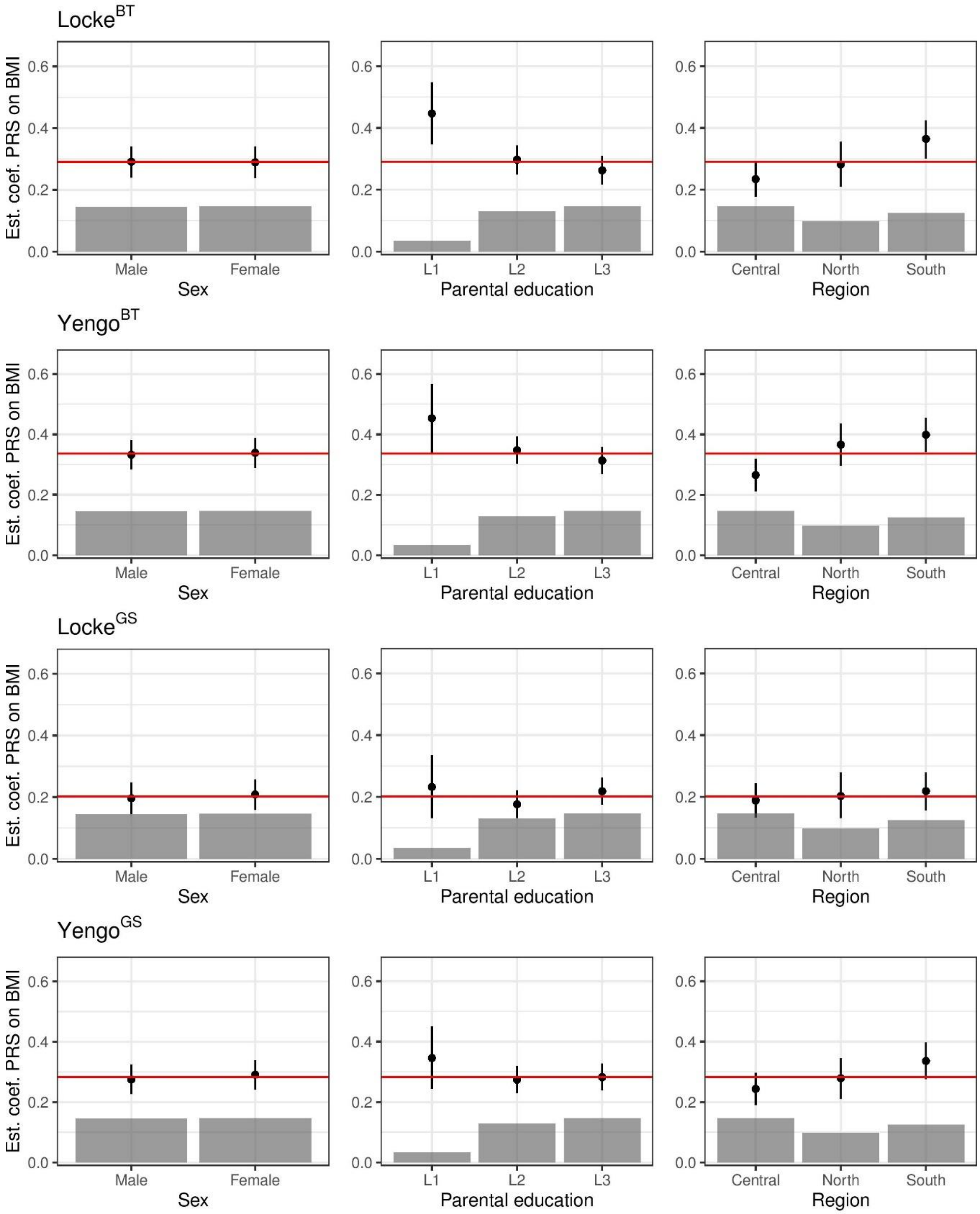


**Figure S5. Interactions of PRS-Locke$^{BT}$, PRS-Yengo$^{BT}$, PRS-Locke$^{GS}$ and PRS-Yengo$^{GS}$ with sociodemographic factors for BMI.** Associations between PRS and obesity are shown in different strata (beta estimates and 95% confidence intervals) as well as in the whole study population (red line). The distributions of the demographic factors are shown in histograms.

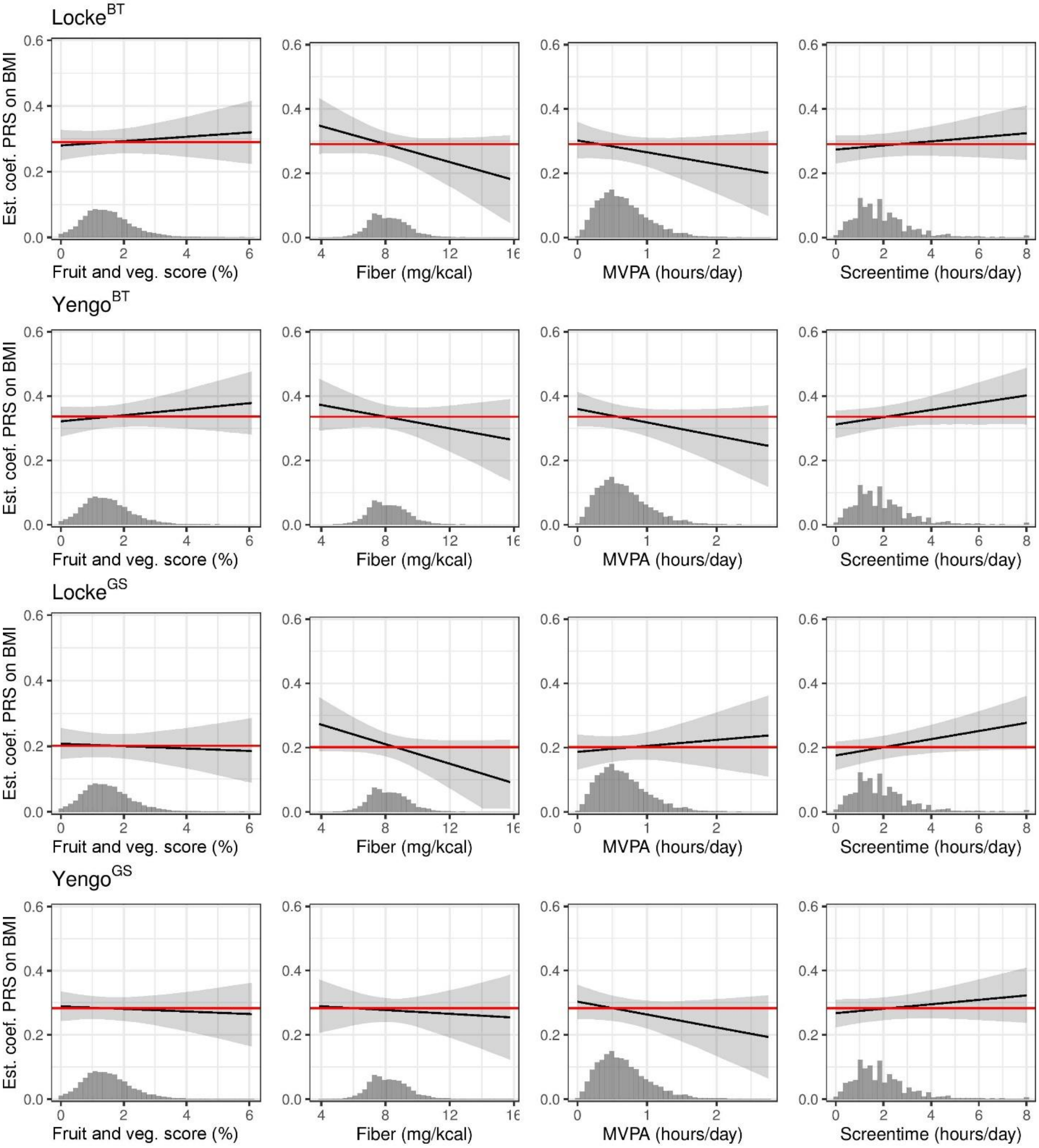


**Figure S6. Interaction of PRS-Locke$^{BT}$, PRS-Yengo$^{BT}$, PRS-Locke$^{GS}$ and PRS-Yengo$^{GS}$ with lifestyle factors for BMI.** Associations between PRS and obesity are shown in dependence of the PRS (beta estimates and 95% confidence intervals) as well as in the whole study population (red line). The distributions of the lifestyle factors are shown in histograms.
